# Widespread data leakage inflates accuracy and corrupts biomarker discovery in cancer drug response prediction

**DOI:** 10.64898/2026.02.05.704016

**Authors:** Amir Asiaee, Jared Strauch, Leila Azinfar, Samhita Pal, Heather H. Pua, James P. Long, Kevin R. Coombes

## Abstract

Drug response prediction models guide biomarker discovery, yet their reported accuracy depends on rigorous cross-validation (CV). We show that supervised feature screening applied to all samples before CV, a widespread practice, introduces data leakage that systematically underestimates prediction error. Across 265 drugs and 1,462 cancer cell lines, leakage-free CV raises mean squared error by 16.6% on average, with near-zero feature overlap between leaked and corrected pipelines (mean Jaccard 0.18; 36.2% of drugs share no features). Despite selecting five times more features, the leaked pipeline recovers known drug targets at nearly the same rate as the corrected pipeline, indicating that the inflated feature sets capture statistical artifacts rather than biological signal. A code-level audit of 32 published methods (2017–2024) confirms leakage in 23 (72%), spanning five distinct modes and cited over 3,000 times. The magnitude of inflation from this single leakage mode is comparable to improvement margins typically claimed over elastic net baselines, raising the possibility that some reported advances reflect evaluation artifacts. We provide a leakage taxonomy, audit guide and reference implementation for leakage-free evaluation.

---

Genomic markers that predict drug sensitivity in cancer cells are routinely nominated from large-scale pharmacogenomic screens such as the Cancer Cell Line Encyclopedia (CCLE) and Genomics of Drug Sensitivity in Cancer (GDSC) [1–4]. Cross-validation (CV) is the standard approach for estimating prediction performance in these high-dimensional settings. However, subtle errors in CV implementation can introduce data leakage—the inadvertent use of test-set information during model training—leading to optimistic performance estimates and unreliable feature rankings [5–8]. Here we quantify one prevalent form of leakage, supervised feature screening performed on the full dataset before CV splits, and document its widespread occurrence across the drug response prediction literature.

We compared two elastic-net pipelines [9, 10] using GDSC drug sensitivity for 265 compounds matched to CCLE molecular features for 1,462 cell lines spanning 31 tissue lineages: (i) an *incorrect* pipeline that applies variance filtering, correlation screening and scaling to the full dataset before CV—a “screen-then-validate” pattern used in the seminal CCLE and GDSC studies [1, 2] that implicitly exposes test-fold labels to the feature selection step, violating the assumption that CV folds are held out [5]; and (ii) a *leakage-free* pipeline that repeats these steps within each fold using training data only (Fig. 1b; Online Methods).

**Fig. 1.**
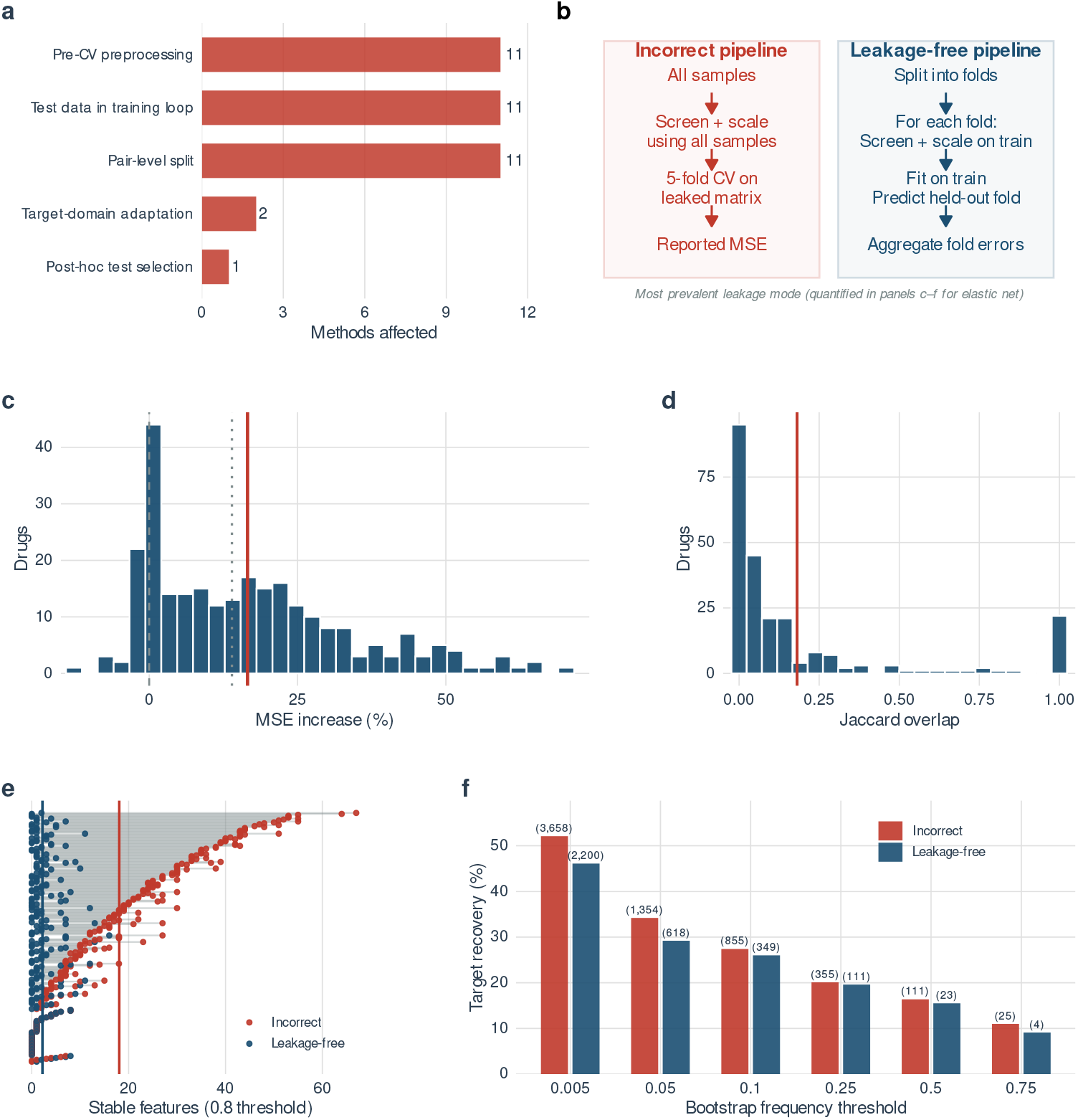
Data leakage inflates performance estimates and destabilizes feature sets across 265 drugs. **a**, Leakage mode distribution across 23 confirmed-leaky methods (Table 1); methods may exhibit multiple modes. **b**, Schematic contrasting the incorrect pipeline (supervised screening on all samples before CV) with the leakage-free pipeline (screening repeated within each fold); this is the most prevalent leakage mode, quantified in panels c–f. **c**, Distribution of MSE inflation (%) under leakage-free CV; solid line, mean (16.6%); dotted line, median (14.0%). **d**, Jaccard overlap between stable feature sets ( ≥ 80% bootstrap threshold); solid line, mean (0.18); 36.2% of drugs have zero overlap. **e**, Stable feature counts per drug under incorrect (red) and leakage-free (blue) pipelines; drugs are ranked by the gap between incorrect and correct feature counts; vertical lines mark means (18.1 vs 2.2). **f**, Drug target recovery rates across bootstrap frequency thresholds for incorrect versus leakage-free pipelines (219 drugs with known targets); numbers in parentheses show mean selected feature counts per pipeline; the incorrect pipeline selects 5× more features yet recovers only marginally more targets (e.g., 16.4% vs 15.5% at the 0.50 threshold), indicating that leakage inflates feature counts without proportionate gains in biological signal.

Leakage-free CV increased the estimated prediction error for most compounds. Across 265 drugs, the corrected pipeline raised mean MSE by 16.6% (median 14.0%), and 83.0% of drugs showed inflated performance under the leaked pipeline (Fig. 1c). The effect was heterogeneous: 35.8% of drugs showed ≥ 20% inflation, 10.9% showed ≥ 40%, and the maximum reached 70.3%. Leakage also destabilized biomarker discovery. The Jaccard overlap between stable feature sets (features selected in ≥ 80% of bootstrap resamples) was low: mean 0.181, median 0.044, with 36.2% of drugs having zero overlap (Fig. 1d). The leaked pipeline produced much larger signatures—on average 18.1 stable features versus 2.2 under the correct pipeline (Fig. 1e). Critically, the inflated feature sets did not translate into proportionately better biological signal: across 219 drugs with known gene targets, the incorrect pipeline recovered any target for only 16.4% of drugs despite selecting five times more features, compared with 15.5% under the correct pipeline (Fig. 1f). These results indicate that leakage simultaneously inflates accuracy and overstates confidence in derived biomarker lists. To assess how broadly leakage extends beyond elastic net, we audited 32 drug response prediction methods spanning classical machine learning and deep learning, published between 2017 and 2024 (Table 1). We categorized leakage into five recurring modes (Supplementary Note 1): (1) preprocessing on all samples before CV, (2) test data used for early stopping or model selection, (3) pair-level splits inconsistent with cold-start claims, (4) target-domain adaptation using test samples, and (5) post-hoc selection of the best test metric. Across the audited methods, 23 of 32 (72%) contained confirmed leakage, three could not be verified due to incomplete public code, and six appeared clean (Table 1; Fig. 1a). The 23 confirmed-leaky methods have accumulated over 3,000 citations (OpenAlex, February 2026). Pre-CV preprocessing was the most prevalent mode, but several methods exhibited multiple leakage types simultaneously. Of the 20 leaky methods for which we could extract a numeric performance margin (Supplementary Table 2), only two directly benchmark against elastic net: DrugCell (5.7% improvement) and TRANSACT (10.4%), both well within our mean leakage inflation of 16.6%. Across the broader set—which uses diverse baselines, metrics and evaluation protocols—the mean claimed improvement is 13.1% (median 9.2%), and 75% claim gains no larger than 16.6%. Because our inflation estimate derives from elastic net, these cross-baseline comparisons are approximate, but the order of magnitude suggests that evaluation artifacts may account for a substantial fraction of reported advances in drug response prediction. Analogous findings reinforce this concern: correcting data leakage in civil war prediction [11], neuroimaging [12] and connectomics [13] eliminated reported advantages of complex models. More broadly, rigorous re-evaluation of train/test methodology has shown that deep learning gene perturbation models do not outperform linear baselines [14] and that drug features contribute zero performance in deep response models [15]. Although we quantify inflation for elastic net, the leakage pattern—applying supervised screening to all samples before any CV split—biases any downstream model [5, 12], and our audit confirms its presence across both classical and deep learning architectures (Table 1).

**Table 1:**
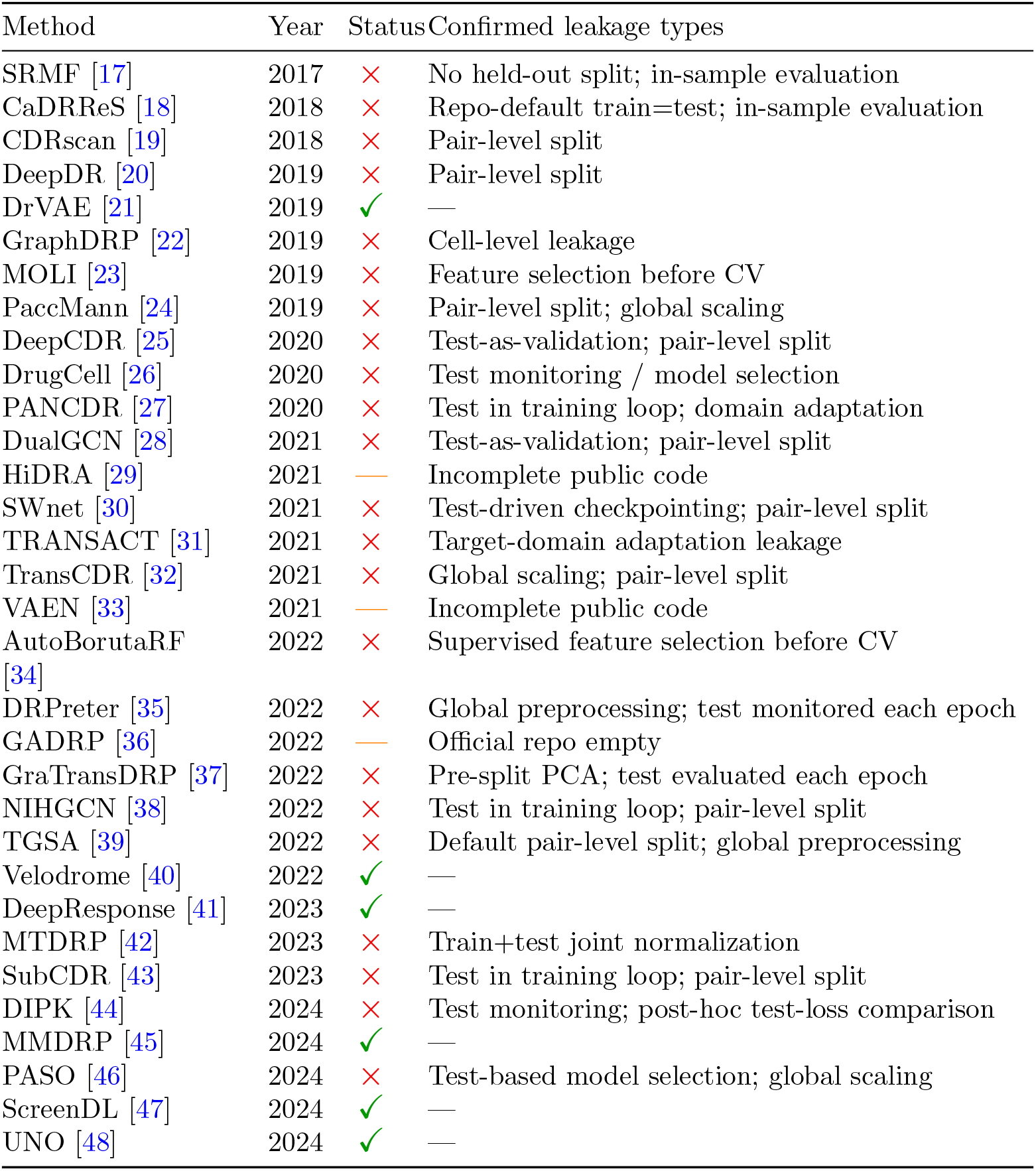
Code-level audit of 32 drug response prediction methods (2017–2024). Status reflects direct inspection of released source code: 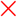 confirmed leakage; 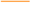 not independently verifiable (incomplete public code); 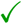 no leakage found. The accompanying code repository provides specific commit hashes, file paths and line numbers for every finding.

Our findings do not imply that prior biological conclusions are necessarily invalid, but that many published performance numbers and feature rankings are likely overconfident. The consequences for biomarker discovery may be particularly severe: leakage inflates stable feature counts by nearly an order of magnitude (Fig. 1e) while providing no proportionate gain in drug target recovery (Fig. 1f), indicating that feature rankings from leaked pipelines are dominated by statistical artifacts rather than biological signal. This undermines downstream applications—drug repurposing, mechanism-of-action studies, patient stratification—where inflated feature lists can misdirect experimental follow-up and biological interpretation. To support improved practice, we provide three concrete deliverables: (1) a five-mode leakage taxonomy with code-level evidence for each audited method, (2) an audit checklist for identifying leakage in new methods, and (3) a reference implementation of leakage-free CV for drug response prediction, available at https://github.com/AsiaeeLab/drug-response-leakage. An independent benchmark effort, DrEval [16], has recently proposed standardized evaluation protocols that complement these resources.

## 1 Online Methods

### 1.1 Data sources

We analyzed drug sensitivity measurements from GDSC/Sanger release 6.0 and molecular features derived from CCLE/DepMap releases distributed with the CCLE update by Ghandi et al. [2–4]. The predictor matrix contained 86,546 features spanning gene expression, copy number, mutation and protein measurements for 1,462 cell lines, and drug response covered 265 compounds (AUC/activity-area values). Data provenance is documented in the accompanying code repository.

### 1.2 Correct and incorrect cross-validation pipelines

For each drug, we fit elastic-net regression models using glmnet [10] to predict response from molecular features. The **incorrect** pipeline applied the full preprocessing stack once using all samples: (i) variance filtering, (ii) correlation screening with the response, (iii) duplicate-feature removal, and (iv) scaling (excluding binary mutation features), and then performed 5-fold CV on the filtered matrix. The **leakage-free** pipeline repeated the same steps independently within each fold, fitting preprocessing on the training split and applying it to the held-out split.

Both approaches used the same hyperparameter grid over the elastic-net mixing parameter (*α*) and selected *λ* by minimum CV error along the regularization path. We report mean squared error (MSE) on held-out folds and compute standard errors as the fold-to-fold standard deviation divided by 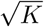, following the standard definition of SE for the mean of *K* fold-level estimates.

### 1.3 Bootstrap feature selection

To quantify feature selection characteristics, we performed bootstrap resampling (200 replicates) and counted how often each feature received a non-zero coefficient. We report counts of selected features, distributions of selection frequencies, and overlap metrics (Jaccard index) at multiple bootstrap frequency thresholds.

### 1.4 Error bar calculation

We report the standard error of the mean 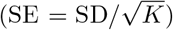 of fold-level errors as the uncertainty measure for CV performance, following the standard definition for the mean of *K* fold-level estimates.

### 1.5 Code-level audit methodology

We audited 32 drug response prediction methods published between 2017 and 2024. Methods were identified through literature searches on PubMed and Google Scholar using terms related to drug response prediction, cell line sensitivity and pharmacogenomics, filtered to those with publicly available source code repositories on GitHub or similar platforms. The selection covers the major model families in the field (matrix factorization, random forests, deep neural networks, graph neural networks and variational autoencoders) but is not exhaustive. For each method, we identified the training and evaluation scripts, traced data flow through preprocessing, splitting and model fitting stages, and classified any leakage patterns into the five-mode taxonomy (Supplementary Note 1). Specific commit hashes, file paths and line numbers supporting each assessment are documented in the accompanying audit guide.

### 1.6 Use of large language models

Large language models were used to assist with drafting and editing the manuscript text and with code-level auditing. All scientific claims, quantitative results and classifications were verified against the repository source code and result files by the authors, who take full responsibility for the content.

### 1.7 Reporting summary

No human participants or animal experiments were involved.

## Data availability

Processed result tables and figures are available in the accompanying repository at https://github.com/AsiaeeLab/drug-response-leakage. The underlying CCLE/DepMap molecular data and GDSC drug response data are available from their original sources.

## Code availability

All analysis code, including leakage-free CV implementations, figure reproduction scripts and the code-level audit guide, is available at https://github.com/AsiaeeLab/drug-response-leakage.

## Author contributions

A.A. conceived the study, discovered the leakage pattern, designed the evaluation framework, implemented the analysis pipeline, conducted the code-level audit and wrote the manuscript. J.S. contributed to discovering the leakage in the original CCLE study and to data processing and pipeline implementation. L.A. crafted parts of the manuscript. S.P. edited the manuscript. H.H.P. provided biological interpretation and edited the manuscript. J.P.L. contributed statistical methodology guidance and edited the manuscript. K.R.C. supervised early stages of the project and edited the manuscript.

## Competing interests

The authors declare no competing interests.

## Supplementary Information

### 1 Supplementary Note 1 — Leakage taxonomy

We identified five recurring modes of data leakage in drug response prediction methods. This taxonomy was developed through code-level inspection of 32 published methods and is intended as a practical audit guide.

1. **Pre-CV preprocessing leakage**: normalization, PCA, variance filtering or supervised feature selection fit on all samples before cross-validation. This is the most prevalent mode (Fig. 1a in the main text) and the one we quantify experimentally. When screening statistics (e.g., correlation with the response) are computed on the full dataset, test-fold labels influence which features enter the model, inflating apparent accuracy and producing non-replicable feature sets.
2. **Test set in training loop**: the test or validation split is used for early stopping, model selection or hyperparameter tuning during training. Even when a nominal train/test split exists, monitoring test-set performance across epochs and selecting the best checkpoint introduces leakage that can bias final reported metrics.
3. **Pair-level split mismatch**: random splits on (cell line, drug) response pairs when the stated goal is generalization to new cell lines or new drugs. Because the same cell line (or drug) can appear in both training and test sets under pair-level splitting, the model may learn cell-line-specific or drug-specific patterns rather than generalizable drug–genomic relationships.
4. **Target-domain adaptation leakage**: domain adaptation or transfer learning methods that use unlabeled test-domain samples during training. When the test domain is the target domain and its samples (even without labels) are used to learn domain-invariant representations, information about the test distribution leaks into the model.
5. **Post-hoc test selection**: reporting the best test metric across epochs, hyperparameter configurations or random seeds without a separate held-out evaluation. This optimistic selection can inflate reported performance even when the training procedure itself is leak-free.

The full audit table is presented as Table 1 in the main text. The accompanying code repository includes direct links to specific commits, file paths and line numbers supporting each assessment.

### 2 Supplementary Note 2 — Additional methodological details

#### 2.1 Variance filtering, supervised screening and scaling

For elastic net baselines, preprocessing consisted of:

- **Variance filter**: retain features with variance *>* 0.01.
- **Correlation screen**: retain features with |cor(feature, response) | *>* 0.1.
- **Duplicate removal**: remove duplicated columns after filtering.
- **Scaling**: z-score continuous features using training-set mean and standard deviation; binary mutation features (identified by the suffix pattern <monospace>mut.</monospace>) were excluded from scaling.

In the incorrect pipeline, these steps were computed once using all samples (leakage). In the leakage-free pipeline, they were recomputed within each fold using training data only. Reference implementations are provided in the accompanying code repository.

#### 2.2 Hyperparameter selection

Both pipelines used the same grid over the elastic-net mixing parameter *α* ∈ *{*0.1, 0.2, 0.3, 0.4, 0.5, 0.6, 0.7, 0.8, 0.9, 1.0*}* and selected *λ* by minimum 5-fold CV error along the regularization path. The hyperparameter combination yielding the lowest mean CV MSE was selected for each drug.

### 3 Supplementary Table 1 — Threshold sensitivity analysis

Jaccard and stable feature counts across bootstrap frequency thresholds. (incorrect vs leakage-free CV pipeline):

3 Supplementary Table 1 — Threshold sensitivity analysis **Jaccard and stable feature counts across bootstrap frequency thresholds** (incorrect vs leakage-free CV pipeline):
| Threshold | Mean Jacc. | Median Jacc. | Zero overlap (%) | Features (incorrect) | Features (leak-free) |
| --- | --- | --- | --- | --- | --- |
| 0.50 | 0.147 | 0.103 | 16.5 | 68.6 | 9.8 |
| 0.60 | 0.128 | 0.071 | 21.3 | 49.4 | 5.9 |
| 0.70 | 0.107 | 0.053 | 28.0 | 33.9 | 3.2 |
| 0.80 | 0.094 | 0.034 | 42.9 | 21.5 | 1.7 |
| 0.90 | 0.091 | 0.000 | 65.8 | 10.9 | 0.8 |

Values derived from 265 drugs; stable features defined as those selected in ≥ threshold fraction of 200 bootstrap resamples. Across all thresholds, the incorrect pipeline selects substantially more features with low overlap to the leakage-free pipeline.

### 4 Supplementary Table 2 — Margin-of-leakage comparison

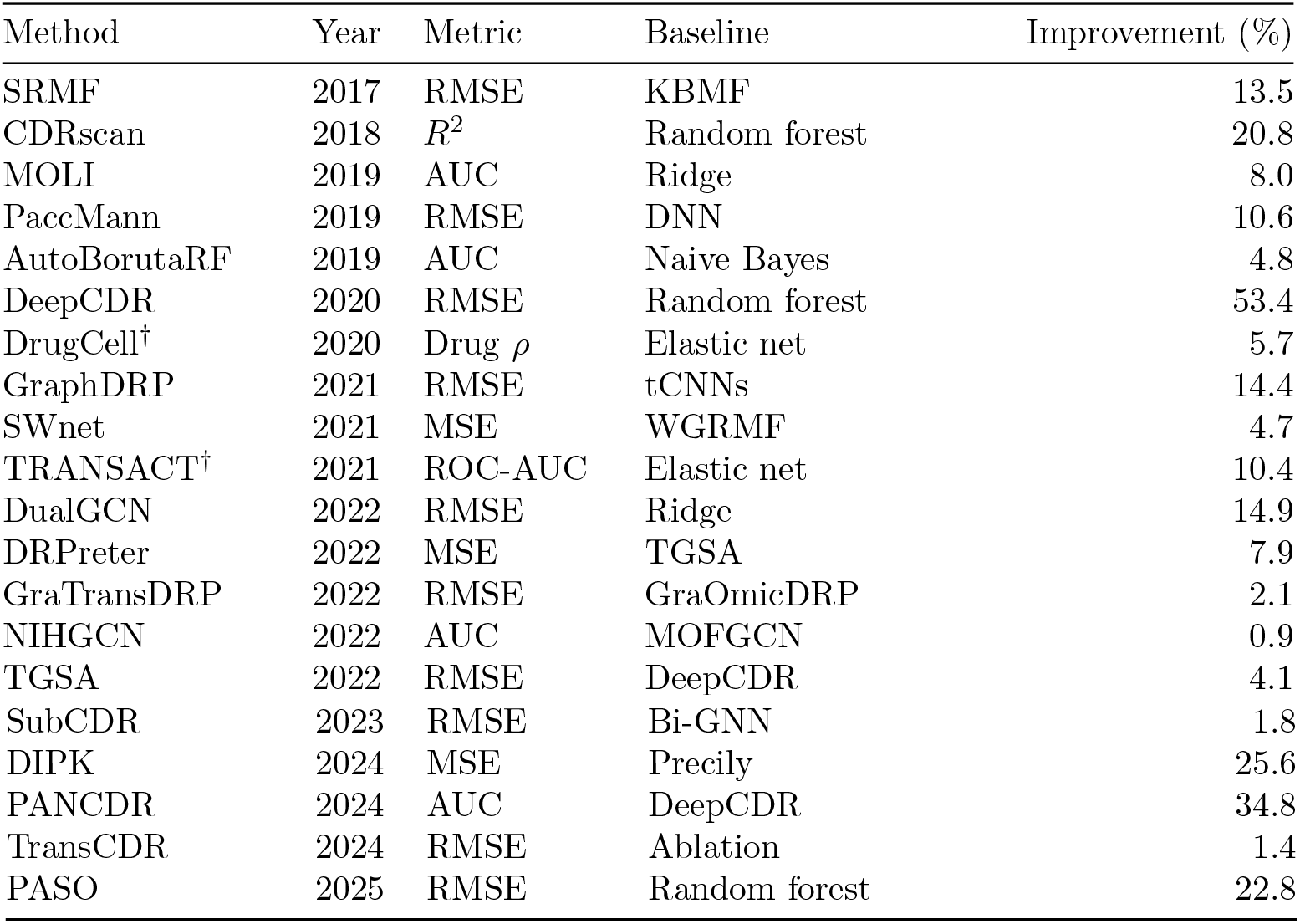
Claimed performance margins of leaky methods. For each of the 20 leaky methods with an extractable numeric improvement from the original publication, we report the primary metric, strongest baseline comparator, and claimed relative improvement (%). Rows marked with † compare directly against elastic net—only these two rows permit a like-for-like comparison with our measured leakage inflation (mean 16.6%). All other baselines differ in model class, metric or evaluation protocol, so their improvement percentages serve as order-of-magnitude context rather than direct comparisons to our elastic-net-specific inflation estimate.

| Method | Year | Metric | Baseline | Improvement (%) |
| --- | --- | --- | --- | --- |
| SRMF | 2017 | RMSE | KBMF | 13.5 |
| CDRscan | 2018 | $R^2$ | Random forest | 20.8 |
| MOLI | 2019 | AUC | Ridge | 8.0 |
| PaccMann | 2019 | RMSE | DNN | 10.6 |
| AutoBorutaRF | 2019 | AUC | Naive Bayes | 4.8 |
| DeepCDR | 2020 | RMSE | Random forest | 53.4 |
| DrugCell† | 2020 | Drug $\rho$ | Elastic net | 5.7 |
| GraphDRP | 2021 | RMSE | tCNNs | 14.4 |
| SWnet | 2021 | MSE | WGRMF | 4.7 |
| TRANSACT† | 2021 | ROC-AUC | Elastic net | 10.4 |
| DualGCN | 2022 | RMSE | Ridge | 14.9 |
| DRPreter | 2022 | MSE | TGSA | 7.9 |
| GraTransDRP | 2022 | RMSE | GraOmicDRP | 2.1 |
| NIHGCN | 2022 | AUC | MOFGCN | 0.9 |
| TGSA | 2022 | RMSE | DeepCDR | 4.1 |
| SubCDR | 2023 | RMSE | Bi-GNN | 1.8 |
| DIPK | 2024 | MSE | Precily | 25.6 |
| PANCDR | 2024 | AUC | DeepCDR | 34.8 |
| TransCDR | 2024 | RMSE | Ablation | 1.4 |
| PASO | 2025 | RMSE | Random forest | 22.8 |

### 5 Supplementary Figure 1

**Fig. 1. Supplementary Fig. 1.**
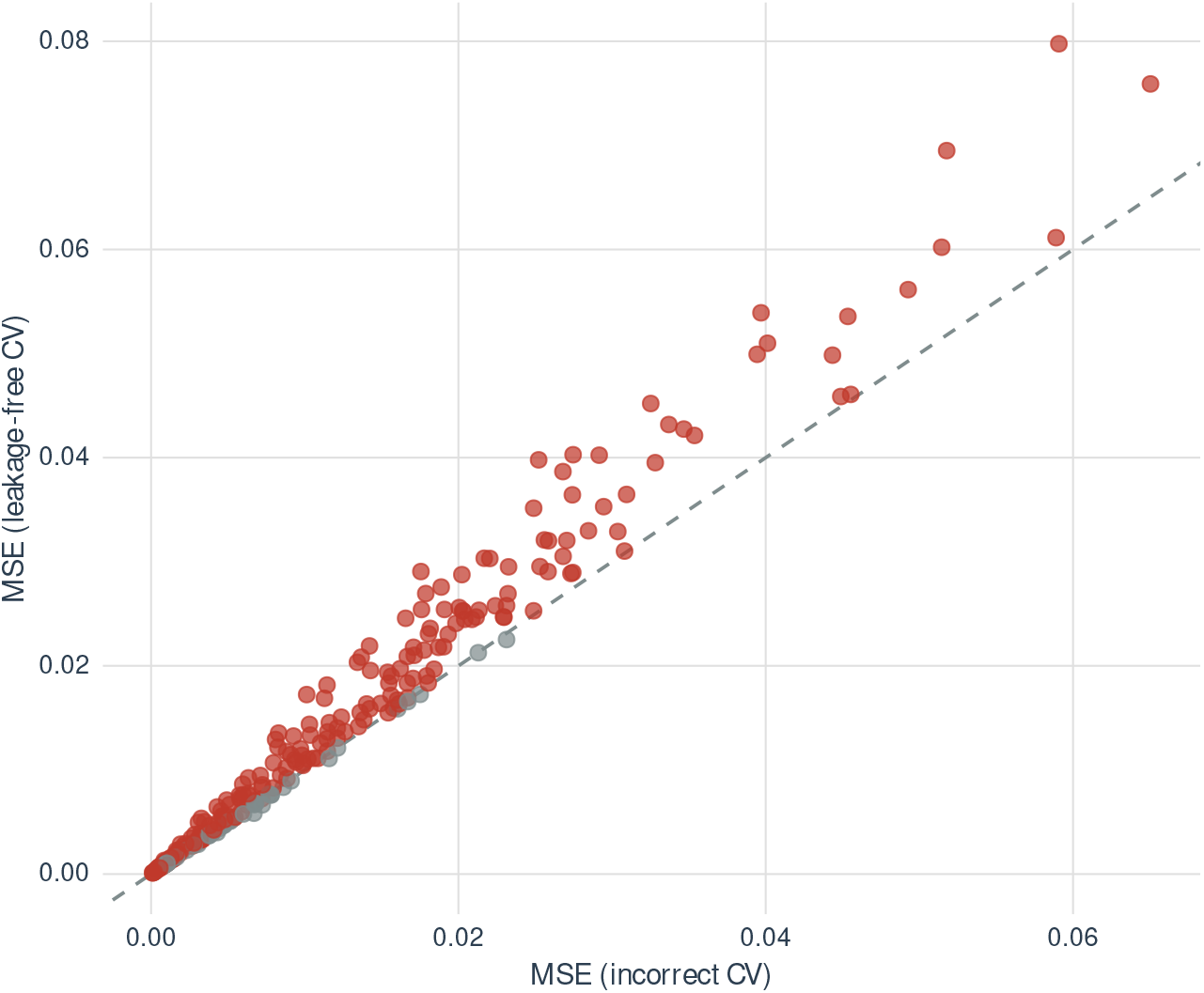
Drug-wise MSE under incorrect versus leakage-free CV. Each point represents one of 265 drugs. Points above the diagonal indicate higher MSE under leakage-free CV (i.e., the incorrect pipeline underestimated error). Of 265 drugs, 83.0% fall above the diagonal. Points are colored by direction: red indicates drugs where leakage-free CV reveals higher MSE; gray indicates drugs with no increase.

### 6 Code and Data Availability

All analysis code is available at https://github.com/AsiaeeLab/drug-response-leakage, including:

- Reference implementations of correct and incorrect CV pipelines
- Complete analysis notebooks reproducing all figures and tables
- Leakage audit guide with code links for each audited method
- Processed result tables and supplementary data files

## Notes

### Competing Interest Statement

The authors have declared no competing interest.

### Summary of Updates

The number of audited methods increased from 12 to 32. The new method (Data Shared Elastic Net) was removed from the paper and the manuscript is now a focused methodological critique. Key additions: (1) a five-mode leakage taxonomy with code-level evidence for all 32 methods, (2) a margin-of-leakage analysis showing that 75% of leaky methods claim improvements smaller than our measured leakage inflation, (3) drug target recovery analysis demonstrating that leaked pipelines select five times more features without proportionate gains in biological signal, and (4) a public code repository with audit guide and reference implementations. The title was updated to reflect the biomarker discovery findings.

